# DeepSlice: rapid fully automatic registration of mouse brain imaging to a volumetric atlas

**DOI:** 10.1101/2022.04.28.489953

**Authors:** Harry Carey, Michael Pegios, Lewis Martin, Christine Saleeba, Anita Turner, Nicholas Everett, Maja Puchades, Jan Bjaalie, Simon McMullan

## Abstract

Registration of data to a common frame of reference is an essential step in the analysis and integration of diverse neuroscientific data modalities. To this end, volumetric brain atlases enable histological datasets to be spatially registered and analysed, yet accurate registration remains expertise-dependent and slow. We have trained a neural network, DeepSlice, to register mouse brain histology to the Allen Brain Atlas, retaining accuracy while improving speed by >1000 fold.

## Main

A core concept in neuroscience is *functional localization*, in which different aspects of the brain’s operation are mediated by physically distinct neural circuits. In this context, a necessary step in the analysis of neural data is their registration to the brain regions from which they were derived. In the mouse (the most widely used species in contemporary experimental neuroscience) the gold-standard tool for registration of neuroanatomy is the Allen Mouse Brain Atlas, which delineates the adult mouse brain into hundreds of structures within a standardized coordinate system, the Allen Common Coordinate Framework (CCF)^1^.

Although long-recognized as a critical step for standardization and integration of complex multimodal datasets^2,3^,registration of data to coordinate systems such as the CCF remains laborious and reliant on human skill^4,5^. Existing tools typically assume programming proficiency, take weeks or even months to register typical datasets, and offer low levels of automaticity ^6-10^. In contrast, Convolutional Neural Networks (CNNs) have shown great promise in the automated analysis of other types of imaging data, including cellular histology^11^ and pose estimation^12^, but to date no equivalent tool for the registration of neuroimaging data to volumetric atlases such as the CCF has been described. Here we detail the development and performance of DeepSlice, a CNN trained on a massive histological dataset for the automatic alignment of coronal mouse brain histology to the CCFv3, which we provide as an open-source Python package (github.com/PolarBean/DeepSlice) and a fully functional online tool (www.DeepSlice.com.au).

DeepSlice is based on the Xception CNN^13^, modified such that the final classification layer was replaced with nine linear output neurons able to regress the corresponding CCF anchoring vectors (O_xyz_, U_xyz_, and V_xyz_) used by the QuickNII histological alignment tool^4^ (Figure 1A). DeepSlice was initially trained (Figure 1B) using a curated dataset of coronal mouse brain sections obtained from the Allen Brain Institute, which had been aligned to the CCF. This included 131k slide-mounted histological sections from the Allen Gene Expression Atlas^14-16^ (AGEA), processed for *in situ* hybridization, immunohistochemistry or Nissl staining and imaged using brightfield or fluorescence optics, and 443k sections from the Allen Connectivity Atlas^17^, which contained multichannel images of viral reporter expression captured using serial 2-photon block-face imaging (S2P). Prototype models of DeepSlice showed good generalizability to different staining and imaging modalities but poor accuracy overall, potentially reflecting shortcomings in the methodology used to register the AGEA datasets used for training, as previously reported^18,19^ and indicated by *post-hoc* analysis of the AGEA dataset (Supplementary Figure 1). We therefore augmented DeepSlice with a second round of training, using 920k synthetic histological images propagated from plausible anchoring coordinates (Supplementary Figure 2) and rendered using the Allen S2P^17,18^ and Nissl^15,16^ template volumes (Figure 1B). We enhanced variability in all training datasets by random addition of noise, pixel drop-out, and warping of input images (Supplementary Figure 3). We found Allen Connectivity Atlas registrations to be more accurate than AGEA registrations, and therefore included them in both phases of training. This approach yielded excellent generalizability and accuracy that extended to unseen imaging modalities. Alignment vectors predicted by DeepSlice corresponded closely with registration metadata from ten unseen Allen Connectivity Atlas experiments (1400 images in total), quantified by estimating the average Euclidean distance (in 25 µm CCF voxels) that any neuron in a given section lay from its Allen-assigned position. For each dataset, median DeepSlice performance was between 5.9 and 9.3 voxels (average 7.7 ± 1.0 voxels, ∼190 μm), with less accuracy observed in sections closer to the rostral and caudal limits of CCF space (Figure 1C & D, data presented as mean ± standard deviation throughout).

**Figure 1.**
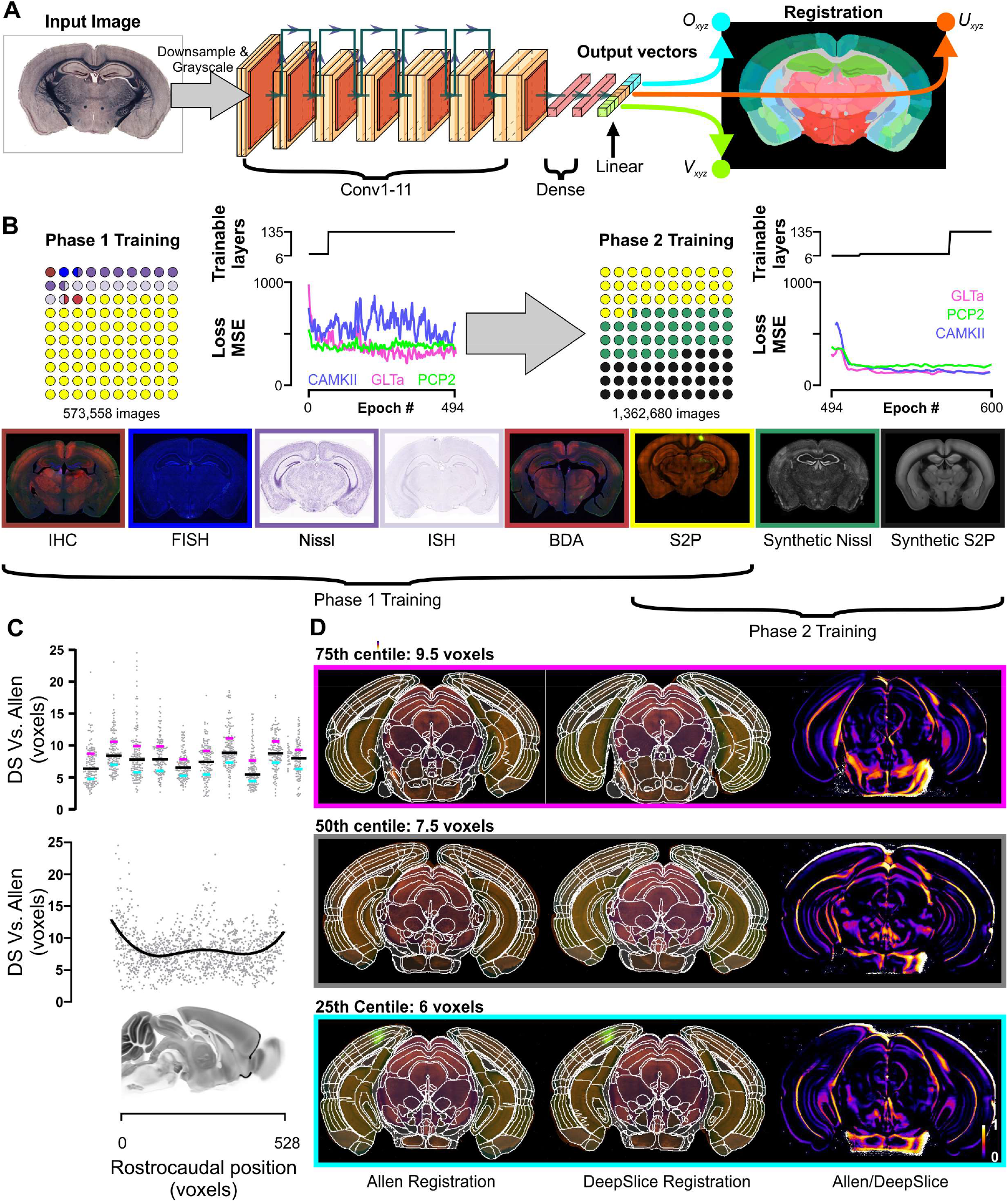
A. DeepSlice is a convolutional neural network that analyses coronal mouse brain images and predicts output vectors (O_xyz_, U_xyz_, and V_xyz_) corresponding to the coordinates of the image corners. B. Training was conducted in two phases; the first used 573k images of slide-mounted and serial 2-photon (S2P) sections, downloaded with their alignment metadata from the Allen Gene Expression (131k) and Connectivity Atlases (443k). The second phase used the same library of S2P images supplemented with a further 0.9M synthetic coronal sections generated through the Allen Nissl and S2P volumes. Loss plots show performance against three human-aligned datasets of slide-mounted histology (n = 119 sections). C. Differences in predicted positions between DeepSlice-and Allen-generated registrations in 1,400 sections from 10 S2P experiments that were omitted from the training set. Lines in upper panel denote median and interquartile ranges; lower panel shows rostrocaudal distribution of error with a sagittal view of the mouse brain for reference. D. Comparison of DeepSlice-generated registrations to corresponding Allen registrations; heat plots highlight differences between registrations. Examples shown are representative of 75^th^, 50^th^ and 25^th^ centile performance across 10 S2P datasets.

The S2P imaging used in the Allen Connectivity Atlas is conspicuous in its uniformity and high quality, and is not representative of the slide-mounted histology used by most contemporary researchers, which is subject to misalignment in dorsoventral and mediolateral cutting angles, tears, bubbles, and deformities. Quantification of DeepSlice performance against slide-mounted sections from the AGEA was inappropriate due to the limitations of the AGEA registration methodology^18^, so we developed a benchmarking dataset consisting of 305 unseen slide-mounted sections from 7 mouse brains^5,16,20-23^ (Supplementary Figure 4), processed using a variety of common staining techniques. These sections were registered by 7 human operators of varying abilities (3 ‘novice’ undergraduate students, 2 ‘intermediate’ postgraduate/postdoctoral researchers, and 2 ‘expert’ neuroanatomists with>10 years experience). We randomly designated three brains as Validation (n=119 sections), used to guide model development, and four as Test (n = 191 sections), on which only the final model was assessed.

To account for inter-rater variability of human operators, benchmarking was quantified with respect to ‘ground truth’ registration coordinates calculated from the collated average of human-generated anchoring vectors (Supplementary Figure 5), harnessing the Wisdom of the Crowd phenomenon in which the average output of a group of individuals of varying abilities may surpass the performance of its most proficient members when error is normally distributed^24,25^. As expected, error (the average Euclidean distance that neurons lay from the ground truth position) was inversely related to expertise, with novices performing worse than intermediate or expert aligners (median error across seven datasets: 11.2±3.1 vs. 10.3±3.8 vs. 7.8±2.2 voxels for novices, intermediates, and experts respectively, Tukey P<0.0001 for all comparisons, Figure 2A). Interestingly, the different datasets also represented a major source of variation in performance irrespective of experience level (2-way ANOVA P=0.0002, F (6, 28) = 6.7), matching participants’ experience that some datasets were easier to align than others and suggesting that the Validation and Test sets were sufficiently diverse to generate meaningful data.

**Figure 2.**
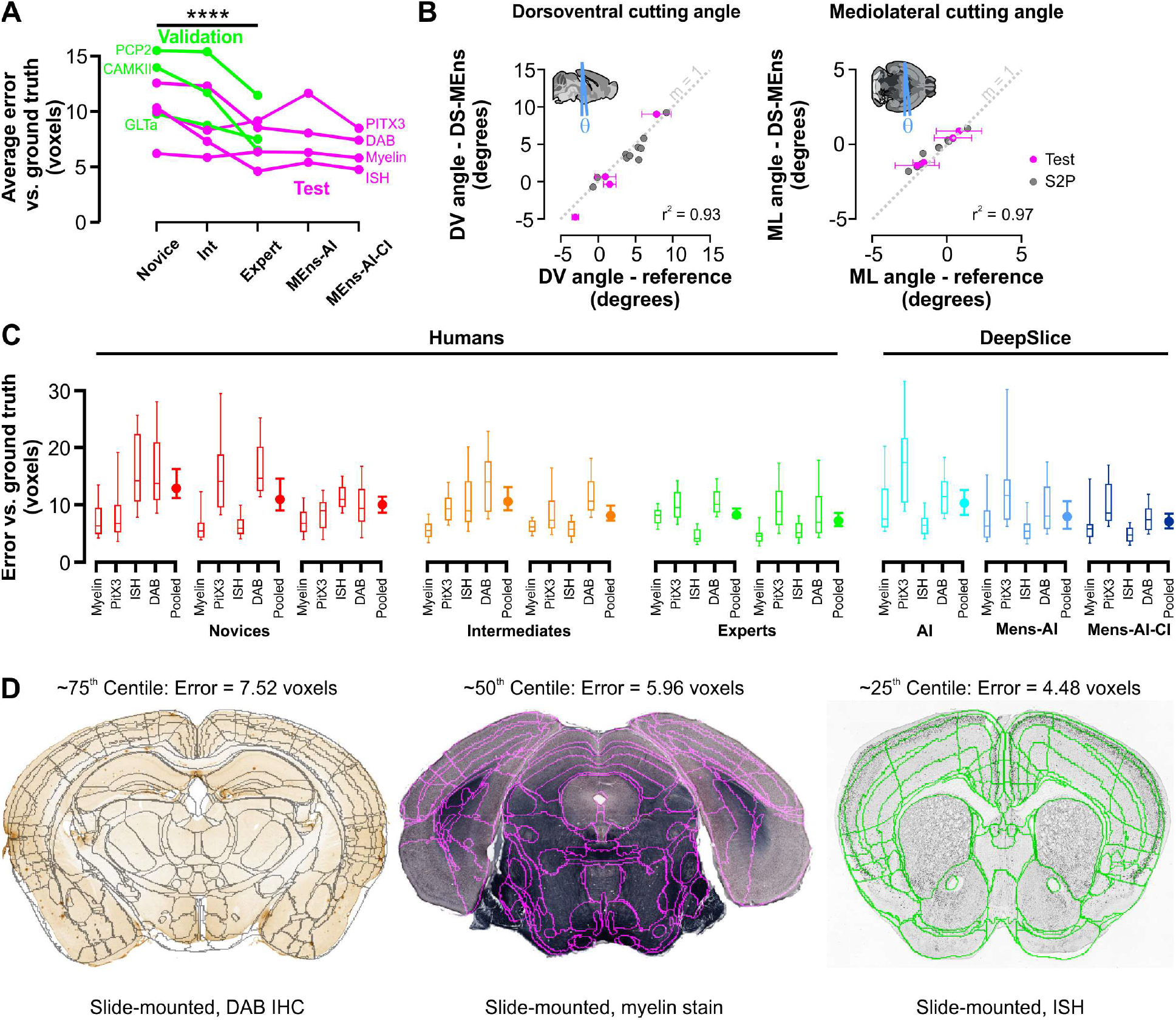
Comparison of DeepSlice vs. human performance on slide-mounted coronal mouse-brain sections. A. Human alignment performance varied significantly by level of expertise on 7 datasets, which were randomly assigned as Validation (green), used for model refinement and development, and Test (magenta), used for benchmarking of final model performance. B. DeepSlice-predicted dorsoventral (left scatterplot) and mediolateral (right scatterplot) cutting angles corresponded closely with cutting angles in benchmark S2P (gray) and human-aligned slide-mounted (black) sections. C. Pooled data show DeepSlice alignment accuracy with reference to human subjects with varying levels of expertise; the ultimate DeepSlice iteration, an ensembled model composed from the top performing 2 CNNs, which integrated cutting index metadata, approximates human expert performance. D. Exemplars from the Test dataset (with overlaid alignment predictions) illustrating approximate overall 75^th^, 50^th^ and 25^th^ centile performance of the Mens-AI-CI model.

DeepSlice initially underperformed on the slide-mounted Test sets compared to hold-out Allen Connectivity Atlas experiments described above, with a median registration error of 10.7 ± 4.7 voxels compared to the human-generated ground truth. This performance was improved by averaging anchoring coordinates generated by the two models which performed best on the Validation sets (Model Ensembling, MEns) and by using cutting index metadata stored in image filenames to weight rostrocaudal position (Cutting Index, CI: Supplementary Figure 6B). Since sequentially cut brain sections share the same dorsoventral and mediolateral cutting angles, we also adjusted the cutting angles of individual predictions such that images from a single experiment reflected the average dorsoventral and mediolateral angles of that dataset, termed Angle Integration (AI). Collectively, these modifications significantly improved DeepSlice performance: predictions of dorsoventral and mediolateral cutting angle were highly correlated with ground-truth measurements in S2P and slide-mounted sections (Figure 2B, r^2^ = 0.93 and 0.97 respectively), and overall alignment accuracy became equivalent to Human Expert level (median error in Test datasets 6.6 ± 1.7 voxels, Figure 2C). These strategies also improved performance on unseen S2P data by a further 25%, significantly reducing the difference between DeepSlice and Allen registrations to 5.8 ± 1.4 voxels (∼140 μm, Tukey P<0.0001).

Although a number of strategies have been developed for the registration of brain histology to reference atlases, to date these require dedicated high powered computing infrastructure, extensive preprocessing of data, a high level of programing literacy and many hours of processing time per brain^9,26^. In contrast, DeepSlice appears to be capable of generalizing across diverse imaging modalities and staining types and does not require preprocessing of images or specialized computer skills, permitting drag-and-drop functionality. DeepSlice is also computationally efficient, even on a mid-range 2019 laptop (74ms/section with a Nvidia GTX 1060 GPU), typically aligning 50-section datasets in under 4 seconds compared to several hours taken by human participants in our study. Such simplicity and efficiency make DeepSlice deployable against enormous neuroimaging datasets.

Furthermore, the .xml output generated by DeepSlice is compatible with existent QuickNII and VisuAlign tools, which are available on the EBRAINS portal (https://ebrains.eu/service/quicknii-and-visualign), making it easy to integrate DeepSlice predictions into established image registration pipelines such as QUINT^5^. Future versions of DeepSlice will be embedded into the EBRAINS online data repository and toolbox, permitting automated alignment of images within a shareable workspace.

DeepSlice does have some weaknesses: like human aligners, DeepSlice relies on visual cues within section to determine position and, like human aligners, tended to underperform on tissue in which grey-white matter contrast is obscured, particularly experiments counterstained with neutral red; it may be that preparations that enhance contrast (such as myelin stain) or closely resemble training data (Nissl and background fluorescence were highly represented in the training sets) improve reliability. Tissue deformity is also likely a source of error; whereas human operators may optimize alignment of deformed sections around the part of the brain that is most relevant to their work, DeepSlice has no such intuition and appears to predict alignments in a more holistic manner. Further warping of training data may make future DeepSlice models more robust to section variability, but such efforts are unlikely to match those delivered by optimal sample preparation.

In conclusion, DeepSlice aligns coronal mouse brain histology to the CCF in a reliable and rapid manner. It represents a means of standardizing and simplifying the analysis of mouse brain histology, while its integration into imaging repositories will facilitate sharing of multimodal neuroanatomical data for independent analysis.

## Figures

**Supplemental Figure 1:**
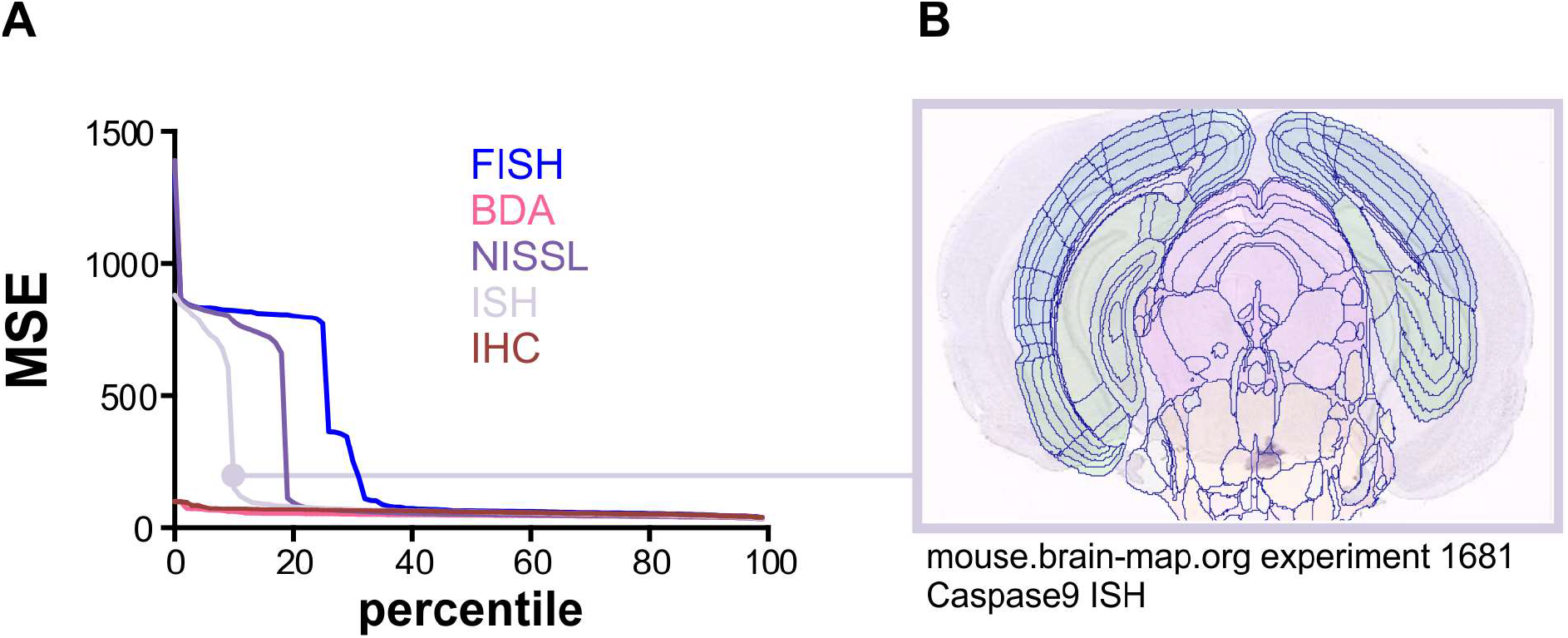
Divergence of prototype DeepSlice predictions from Allen-alignment metadata in FISH, Nissl and ISH datasets (A), which were due to errors in the source data. An example of a poorly aligned Allen experiment is shown in B. MSE in the prototype model was used to filter likely spurious experiments from the training dataset

**Supplementary Figure 2:**
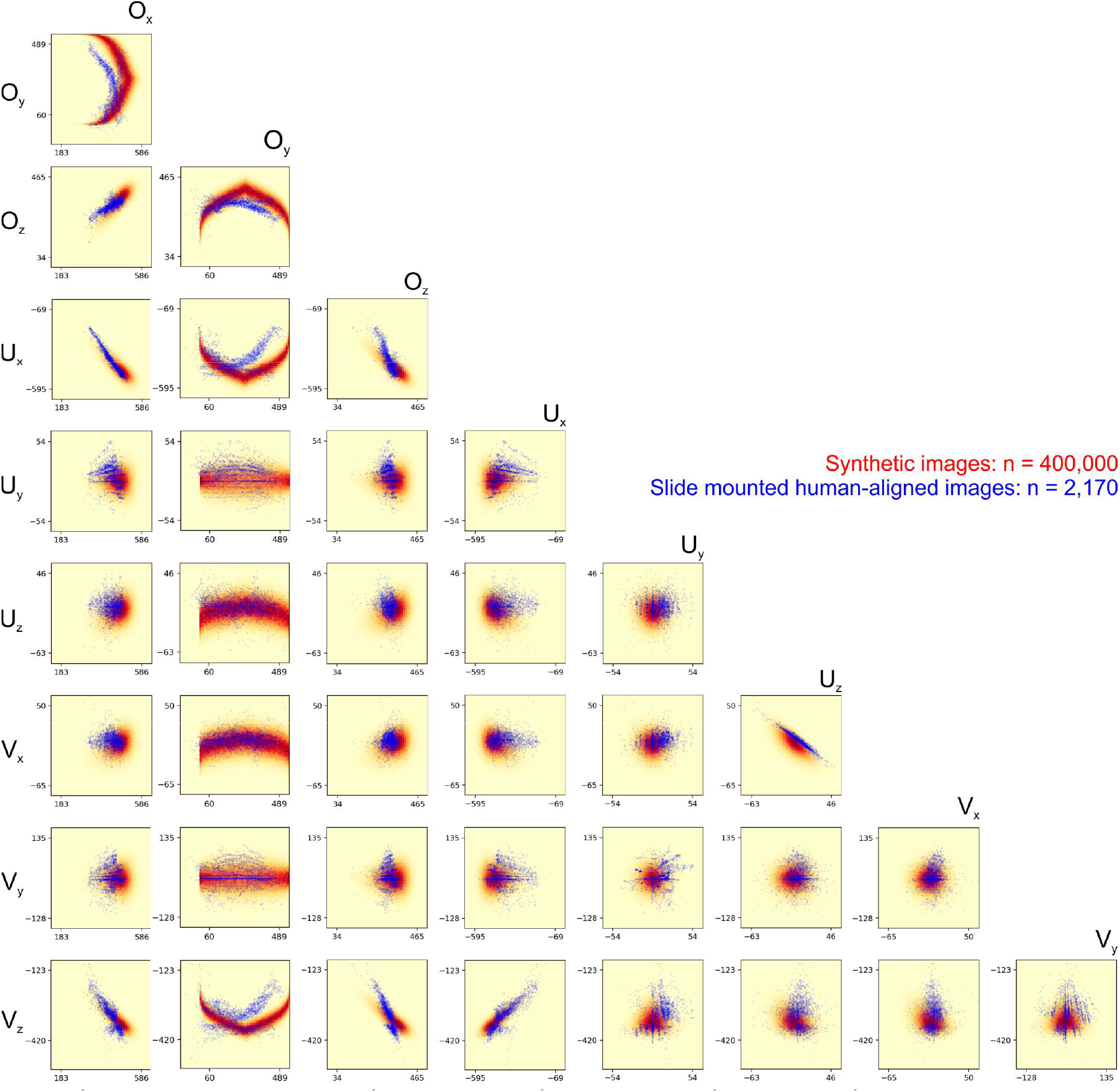
The range and interaction between alignment vectors used to generate 400k synthetic histological images (red) corresponded closely with those assigned by 7 human subjects in the manual alignment of 305 slide-mounted sections (blue).

**Supplementary Figure 3:**
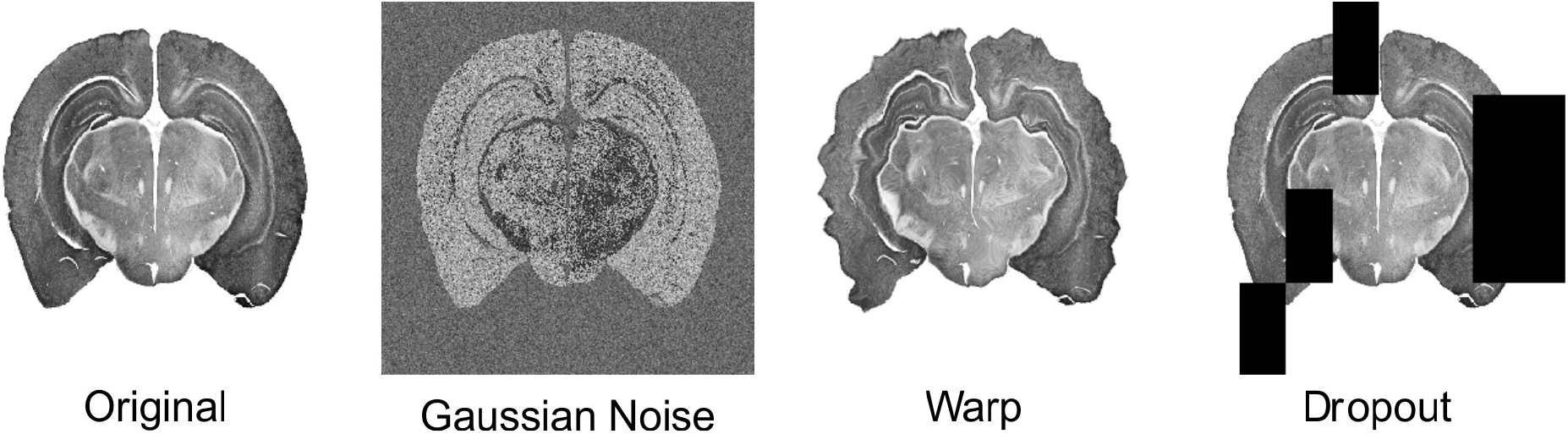
The variability of training data was enhanced by application of filters that randomly assigned combinations of noise, non-linear distortion, and omission of parts of the image. The examples shown illustrate the maximum levels that could be assigned for each filter.

**Supplementary Figure 4:**
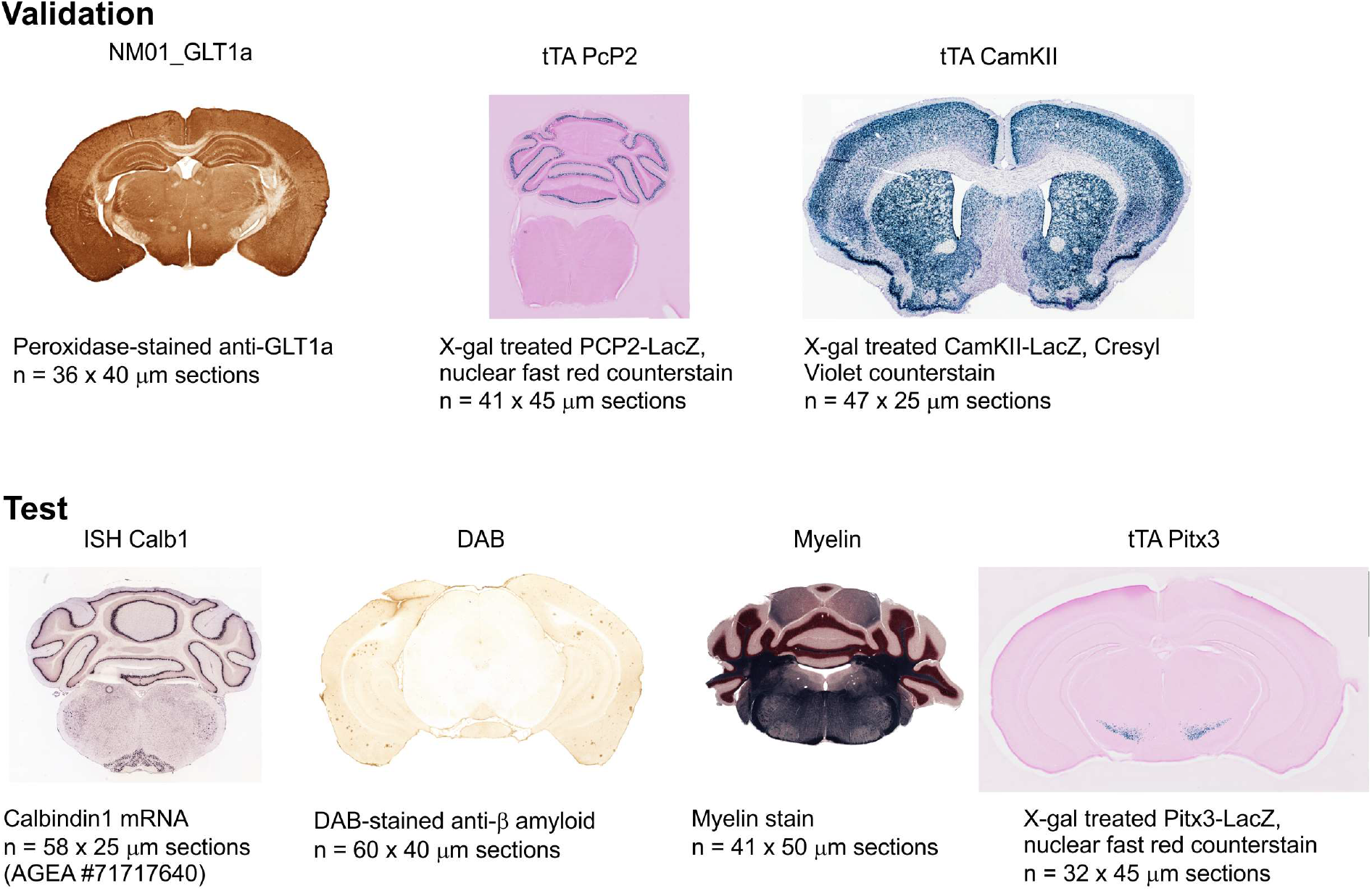
Samples of slide-mounted histology used for generation of Validation^20-22^ and Test^5,16,20,23^ datasets; each dataset was independently aligned by seven human subjects; DeepSlice performance was used to guide model development (Validation) and benchmarking of the final model (Test). The ISH Calb1 dataset was obtained from the Allen Gene Expression Atlas (http://mouse.brain-map.org/experiment/show/71717640) and was not included in the DeepSlice training set.

**Supplementary Figure 5:**
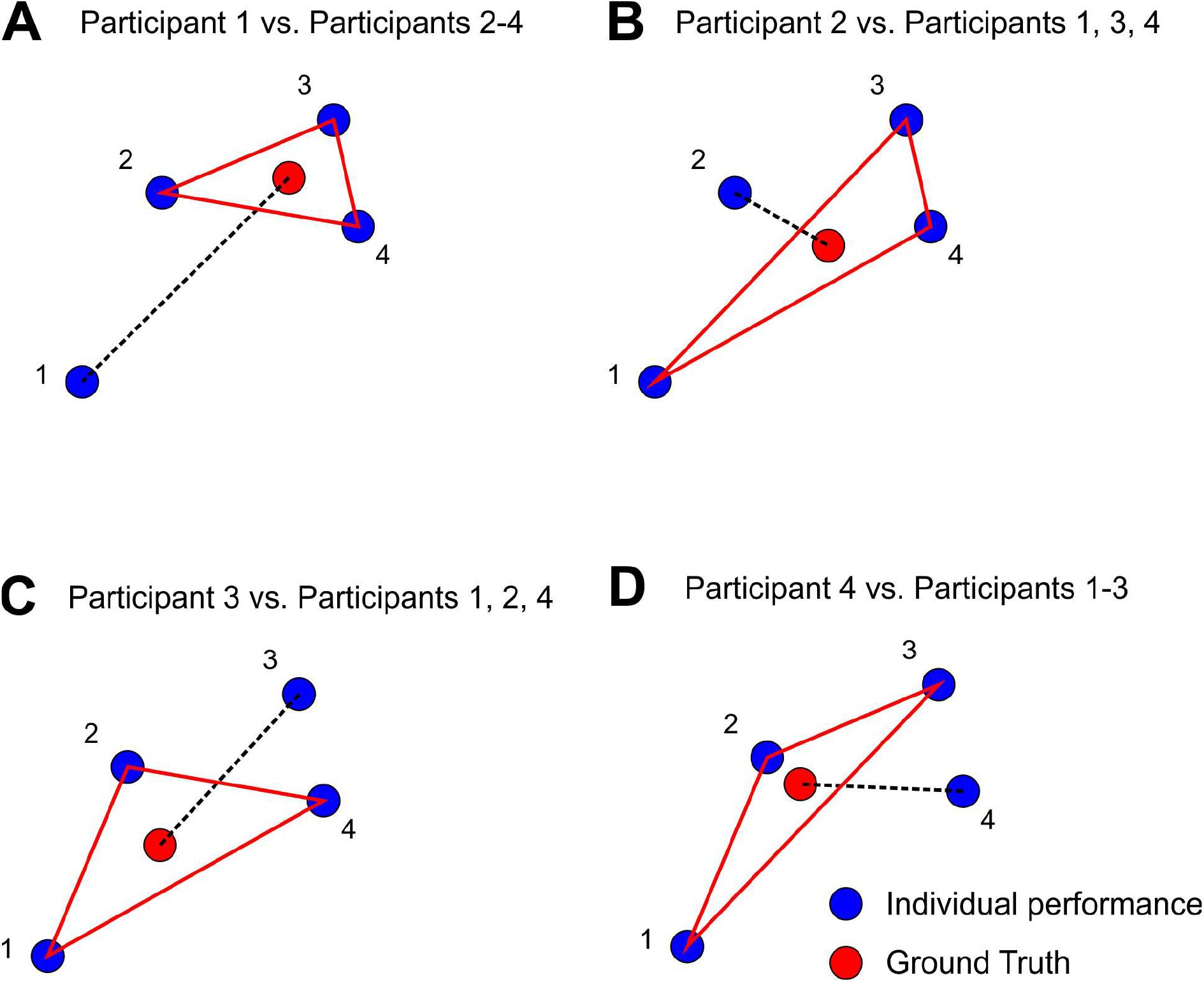
Conceptual overview of approach used for determination of ‘ground truth’ from alignments made by multiple human subjects: anchoring vectors generated by each participant are quantified with respect to the average anchoring vectors produced by the other participants. In this simplified schematic diagram, Participant 2 returns the closest value to the ‘ground truth’ aggregate produced by the other participants (shortest dashed line), whereas Participant 1 returns the furthest value from the ‘ground truth’. A similar approach was used to quantify DeepSlice performance with respect to human operators.

**Supplementary Figure 6:**
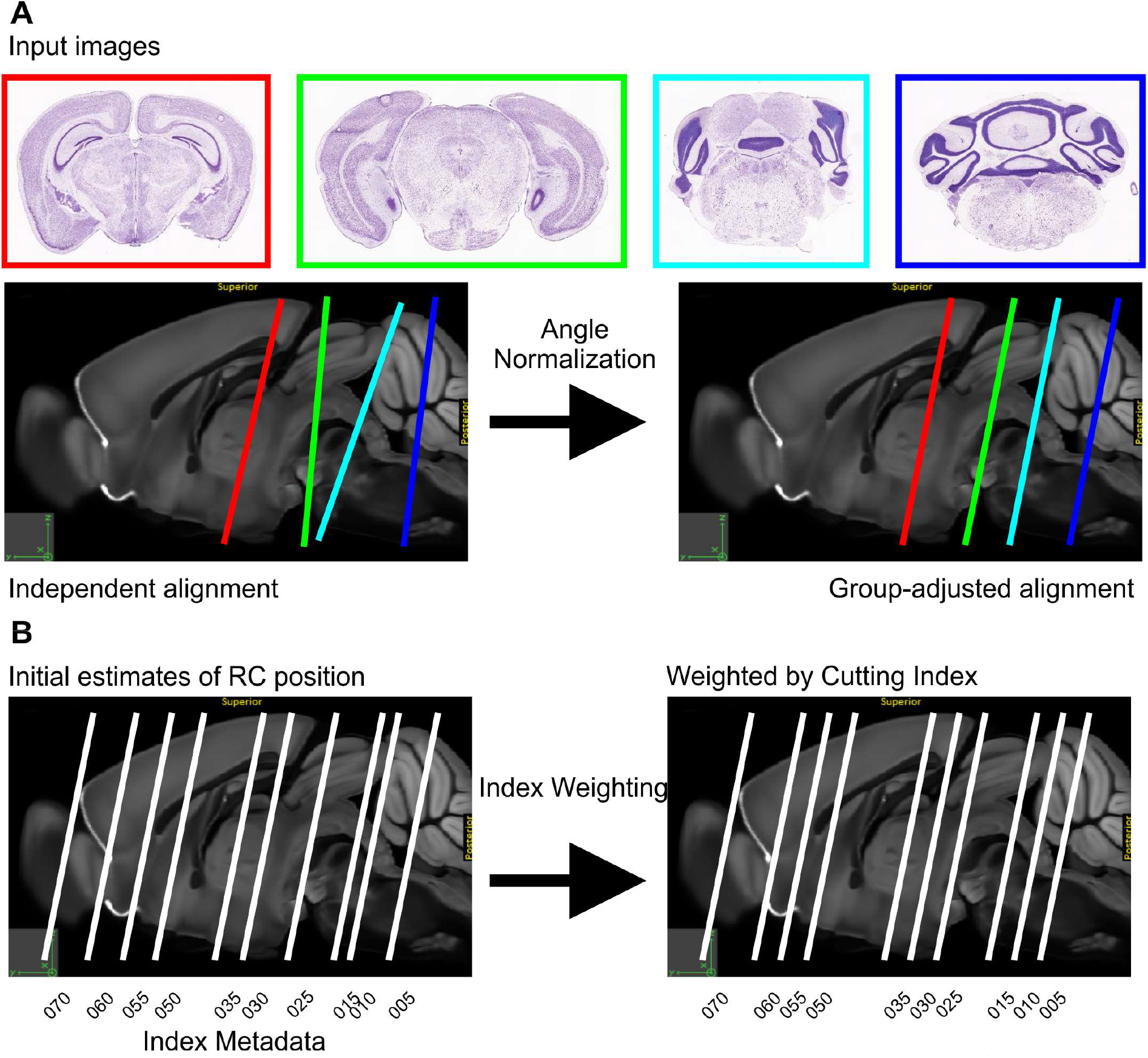
Postprocessing adjustments improved DeepSlice performance; A. Angle Integration (AI) averaged angles made in independent DeepSlice predictions from the same dataset and applied the averaged angle to each image. B. Cutting Index weighting (CI) uses section order metadata to predict the thickness of each section and uses this information to weight rostrocaudal spacing of sections.

## Acknowledgements

Work in the authors’ laboratories is supported by the National Medical & Health Research Foundation, the Hillcrest Foundation, by Macquarie University (SMcM), and from the European Union’s Horizon 2020 Framework Program for Research and Innovation under the Specific Grant Agreement No. 945539 (Human Brain Project SGA3) and the Research Council of Norway under Grant Agreement No. 269774 (INCF, JGB). The authors are grateful to Macquarie University for access to their HPC resources, essential for production of early DeepSlice prototypes. We are grateful to Ann Goodchild for her time-saving blunt assessments of many failed prototypes, for the motivation provided by Dr William Redmond, and especially to Veronica Downs, Freja Warner Van Dijk and Jayme McCutcheon, whose Novice alignments were instrumental to this work. We would like to thank Gergely Csúcs for providing his expertise and many atlasing tools.

## Online Methods

### Model generation and training

Code was written in Python and the model constructed using the Xception architecture, a pretrained CNN consisting of ∼21 million trainable parameters that has high performance on benchmark ImageNet datasets^27^, using the Keras machine learning library (https://keras.io/). The final Softmax layer of Xception was removed and replaced with two dense layers, each consisting of 256 neurons with rectified linear unit activation functions, and nine output neurons with linear activation functions corresponding to the O_xyz_, U_xyz_, and V_xyz_ anchoring vectors used by QuickNII. Xception was initialized with weights pretrained on the ImageNet database^28^. All input images were down-sampled to 299×299 pixels and grayscaled.

Models were optimized using the mean squared error (MSE) loss function and the Adam optimizer^29^, with an initial learning rate of 0.001 and batch size of eight. All Xception layers (except for batch normalization layers) were initially frozen with the only trainable layers being the final dense layers, which were randomly initialized. Layers were iteratively unfrozen as loss plateaued until the entire model was unfrozen (Figure 1B); when the loss plateaued a final time the learning rate was further reduced to 0.0001. Pilot studies revealed that lowering the learning rate beyond 0.0001 yielded no further improvement. Performance against unseen holdout training images and human-aligned slide-mounted Validation sections was plotted every 5000 iterations. Training on an RTX 2080 Ti took 3 to 4 days to reach convergence.

### Image library & curation

Prealigned coronal mouse brain images and corresponding alignment metadata were obtained via the Allen API (http://api.brain-map.org) and converted into O_xyz_, U_xyz_, and V_xyz_ QuickNII anchoring vectors using the Allen2QuickNII script (https://github.com/Neural-Systems-at-UIO/allen2quicknii). During the assessment of prototype DeepSlice models we discovered that many training images returned unexpectedly high MSE values; in such cases, examination of the original data always revealed errors in the original alignment vectors (i.e. contamination of the training dataset: Supplementary Figure 1A). We therefore processed the entire training dataset through a prototype model and sorted images by MSE. For each data type (Nissl, ISH, IHC, FISH, BDA, etc.) we observed an inflection point at which alignment MSE grew exponentially (Supplementary Figure 1B), which was used as a cut-off for elimination of potentially spurious training data. This resulted in a final library of 131,240 curated sections from the Allen Gene Expression Atlas (7,124 IHC, 7,181 FISH, 50,865 Nissl, 57,501 ISH, and 8,207 BDA). No such outliers were identified in 442,680 images from the Allen Connectivity dataset. This library was used to train subsequent DeepSlice models.

### Generation of Synthetic datasets

QSlicer was used to generate a large dataset of virtual sections cut through the Allen Nissl and S2P reference volumes using anchoring coordinates that were representative of those found in real-world images. 520,000 images were propagated using coordinates contained within the Allen Gene Expression Atlas and Allen Connectivity libraries. This data was supplemented with a further 400,000 synthetic images, the coordinates of which were randomly generated based on the Gaussian distribution of anchoring vectors contained within the human benchmark dataset. This resulted in a large dataset of images that closely resembled the dimensions and cutting angles found within human-aligned slide mounted sections (Supplementary Figure 2). Synthetic images that contained minimal brain tissue (e.g. due to tangential cutting angles) were identifiable by small file size (<7 kb, n = 58,000), a result of image compression of their uniform black appearance, and excluded, and a further 20,000 images held out for validation.

### Quantification of performance

Differences between competing alignments of the same image (e.g. DeepSlice vs. Ground Truth) were quantified by calculating the average Euclidean distance of pairs of voxels that correspond to the CCF-projected locations of each pixel in the image, masked such that voxels that fall outside the brain were excluded from analysis. The code used to quantify performance in included in the DeepSlice Github repository (https://github.com/PolarBean/DeepSlice/).

### Estimation of ground-truth

Human alignment of slide-mounted histology is inherently subjective. Such inter-rater variability makes direct comparison of individuals difficult to interpret, as calculated differences quantify the disagreement between human alignments without indicating which one is superior. The same problem applies to comparison of DeepSlice to individual human raters. What is the appropriate ‘ground truth’ when even experts disagree?

One approach is to ask many observers to perform the same task (align the same histological section) and look at the average result. If human error is normally distributed, then the average of the OUV anchoring coordinates likely approximate the ‘true’ position of the section (schematically illustrated in Supplementary Figure 5). To calculate a reliable ‘ground truth’ for evaluation of performance, a group of seven human subjects was trained to use the QuickNII alignment tool. Each subject aligned seven datasets of mouse brain histology, the order of which was counterbalanced to mitigate practice effects. Aligners were classified as either Novice (<1 year of rodent neuroanatomy experience), Intermediate (>2 years’ experience), or Expert (>10 years’ experience). The alignment vectors generated by each human were compared to the arithmetic mean of the alignment vectors generated by the other six human participants (i.e. an alignment corresponding to O_xyz_, U_xyz_, and V_xyz_ vectors generated by Participant 1 was compared to an ensembled alignment calculated by averaging the vectors generated by Participants 2 – 7, and so on). Alignments generated by DeepSlice were compared to ‘Ground Truth’ alignments generated by averaging vectors from all seven human participants.

### Post-processing

#### Normalization of Cutting Angle

Histological sections are iteratively cut in parallel sections from a single block of brain, which is difficult to align perfectly with orthogonal axes and therefore often contains imperfections in cutting angle. Whereas human experts integrate information contained within multiple sections to inform their estimates of dorsoventral and mediolateral cutting angle, DeepSlice analyzes each section independently, leading to section-to-section variability in the predictions of these values. Angle Integration measures cutting angles in every section from within an experiment, averages them, and then adjusts each individual alignment to match the average value of the group (Supplementary Figure 6A).

#### Model Ensembling

We generated multiple DeepSlice models which, although trained using the same combination of images, differed subtly in the numbers of training iterations, level of noise, and the points at which layers were unlocked. Mirroring the ‘wisdom of the crowd’ approach used to establish Ground Truth for human-aligned slide-mounted histology, we also found benefits from averaging alignment vectors produced by the two best DeepSlice models (as determined by performance on the human aligned Validation images), suggesting that sources of error differed in these models and cancelled each other out. Ensembling more than two models was not found to significantly boost performance.

#### Cutting Index

The sectioning index (the order in which histological sections are cut) is often recorded in the filenames of serial histological images. This information can be used to estimate thickness of histological sections after initial alignment by DeepSlice, assuming constant section thickness within a dataset, and subsequently used to adjust predictions of rostrocaudal position (O_y_). Section thickness was calculated by dividing the difference in the value of O_y_ between adjacent sections by the corresponding iterative change in cutting index and averaging this number across the entire dataset.

